# Comprehensive immune profiling of melanoma-draining lymph nodes identifies plasmacytoid dendritic cells as new biomarker for PD-L1 blockade

**DOI:** 10.1101/2025.08.22.671842

**Authors:** Emma Reynaud, Sonya Belherazem, Özge Cicek Sener, Sarah Hofmann, Alperen Acari, Pragati Lodha, Shruthi Hemanna, Nicolò Coianiz, Volker Ast, Sònia Tugues, Lothar C. Dieterich

## Abstract

Immune checkpoint blockade (ICB) has become a powerful weapon in the treatment of melanoma and other cancer types. Still, only a minority of patients profits from this type of therapy. As ICB is associated with considerable adverse effects, predictive biomarkers to select patients for ICB treatment are urgently needed. Here, we comprehensively immunophenotyped primary tumors and tumor-draining lymph nodes (tdLNs) from a panel of ICB-resistant and -sensitive melanoma models in mice. Surprisingly, we found that whereas tumor-infiltrating CD8+ T cells showed strong responses to PD-L1 blockade in ICB-sensitive melanoma, CD8+ T cell profiles in tdLNs were very similar in ICB responder and non-responder models. In contrast, we identified distinct myeloid immune profiles in tdLNs correlating with ICB responsiveness, including a low frequency of activated dendritic cells and a high frequency of plasmacytoid dendritic cells (pDCs). Notably, pDC marker genes and frequencies in tdLNs of melanoma patients correlated with favorable outcome. Thus, the level of pDCs in tdLNs represents a potential new biomarker for ICB in melanoma patients.

## Introduction

The discovery of the CTLA4 and PD-1 immune checkpoints has triggered a revolution in cancer therapy^1,2^. First approved for the treatment of melanoma, a tumor type characterized by a high mutational load and corresponding immunogenicity, antibodies blocking CTLA4 and PD-1 as well as its ligand PD-L1 (CD274) have become a cornerstone in the treatment of multiple cancer types and achieve astonishing outcomes in malignancies previously difficult to treat^3^. However, only a minority of cancer patients treated with immune checkpoint blockers (ICBs) clearly benefits from the therapy. At the same time, ICB therapy is associated with significant risks and adverse effects, in particular the combination of CTLA4 and PD-1 blockade. Therefore, robust biomarkers are urgently needed to stratify cancer patients for ICB therapy.

Among the first biomarkers implemented for ICB therapy were tumor cell-intrinsic parameters such as PD-L1 expression, microsatellite instability, and tumor mutational burden. However, difficulties in standardizing assessment and intra-tumoral heterogeneity have limited the robustness and predictive strength of these biomarkers^4,5^. Furthermore, clinical and experimental evidence has demonstrated that PD-L1 expression by various host-derived cells, including leukocytes and stromal cells, may be equally or even more relevant for immunotherapy outcomes than PD-L1 expression by tumor cells^6–10^. Other ICB biomarkers rely on immunological parameters, such as total CD45^+^ leukocyte infiltration into the tumor microenvironment (TME) or the prevalence of specific tumor-infiltrating lymphocyte (TIL) subsets or activation states^11^. Similarly, circulating leukocytes, including T cells and myeloid subsets such as monocytes and myeloid-derived suppressor cells (MDSCs) have been found to bear predictive value in the context of cancer ICB therapy^12–15^. However, while the robustness of tumor-infiltrating immune parameters suffers from intra-tumoral heterogeneity, circulating leukocyte numbers or activation signatures, although comparably easy to sample and assess, are subjected to non-cancer-related systemic influences and may not always reflect cancer immunity or therapy responsiveness.

Tumor-draining lymph nodes (tdLNs) have a high prognostic value as indicator of tumor dissemination and are an upstaging factor in multiple cancer types including melanoma. Additionally, they have recently moved to the center of attention regarding cancer immunity and ICB. Importantly, tdLNs have been found to harbor a pool of tumor-specific T cells with high proliferative capacity that respond to ICB treatment and rapidly give rise to progenitor exhausted CD8^+^ T cells (Tpex) subsequently infiltrating the TME^16–18^. Furthermore, surgical removal of tdLNs reduces ICB effectiveness in animal cancer models^19^. These findings suggest that ICBs act partially on the LN microenvironment and that LN-infiltrating lymphocytes may be a valuable source for predictive biomarkers towards ICB in patients with an indication for LN biopsy. In comparison, very little is known about myeloid infiltrates in tdLNs and their relation to tumor immunity and ICB outcomes.

Here, we systematically and comprehensively assessed lymphocytic and myeloid immune landscapes in the tumor microenvironment (TME) and in tdLNs in a panel of syngeneic mouse melanoma models, comprising ICB-sensitive and resistant tumors, to map *a priori* differences and immune remodeling upon systemic PD-L1 blockade. Curiously, the CD8+ T cell LN infiltrate was remarkably similar in ICB-sensitive and resistant models and showed reduced exhaustion in response to systemic PD-L1 blockade independently of therapy effectiveness. In contrast, we observed striking differences in the myeloid immune landscape of LNs draining ICB-sensitive and - resistant melanoma. Among those, a high baseline frequency of plasmacytoid dendritic cells (pDCs) correlated with clinical responses in melanoma patients and could thus represent a new predictive biomarker for ICB therapy.

## Results

### The YUMMER1.7 melanoma model is sensitive to PD-L1 blockade

We selected five syngeneic, transplantable mouse melanoma models, B16-F10, HCMel3^20^, the BRAF^V600E^ mutant lines YUMM5.2, YUMM1.7, and YUMMER1.7, a derivate of YUMM1.7 cells generated through UV-induced somatic hypermutation^21,22^. Each cell line was injected intradermally in both flanks of male C57BL/5 mice and screened for ICB sensitivity *in vivo* through treatment with an anti-PD-L1 antibody or a corresponding IgG2b isotype control once the tumor volume had reached 50 mm³. Then, we monitored tumor growth until the first mouse reached a tumor volume of 1 cm³ or a body weight decreased of 20% (Fig. 1A).

**Figure 1.**
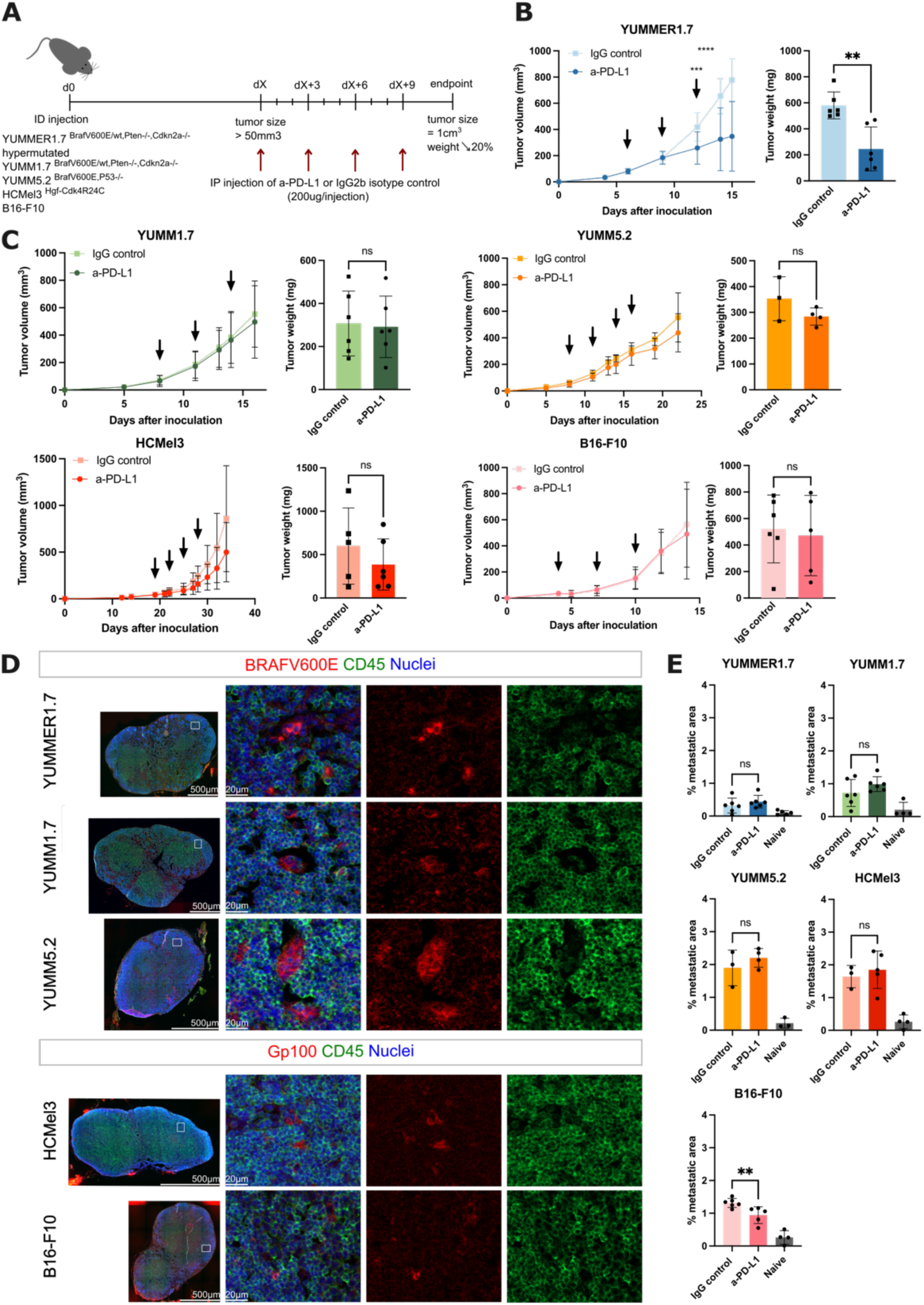
Most transplantable melanoma models do not respond to systemic PD-L1 treatment. **A.** Study design overview. Mice were inoculated intradermally with syngeneic, transplantable melanoma cells and treated with anti-PD-L1 or an IgG2b isotype control when the tumors reached 50 mm³ (as indicated by the arrows). Mice were sacrificed when the tumor volume reached 1 cm³ or the tumor weight decreased by 20%. **B-C**. Tumor growth was monitored by caliper measurement and the tumor weight was recorded at the endpoint. Arrows indicate injection days. N = 3-6 mice/group. ** P ≤ 0.01; *** P ≤ 0.001; **** P ≤ 0.0001; two-way ANOVA with paired Sidak post-test (growth curves) or Student’s t-test (tumor weight); ns=non-significant. **D.** Representative microscopy images of immunofluorescence staining for BRAF^V600E^ or gp100 (red), CD45 (green), and nuclei (blue) of LNs draining the indicated melanoma mouse models treated with the isotype control. **E.** Image-based quantification of metastatic areas in melanoma-draining LNs. N = 3-6 LNs/group. ** P < 0.01; one-way ANOVA with Sidak post-test; ns=non-significant.

Among the 5 models, only the YUMMER1.7 model responded to systemic anti-PD-L1 treatment, showing a significant decrease in tumor volume by day 12 post-inoculation (Fig. 1B). At the endpoint, the tumor weight in the anti-PD-L1-treated YUMMER1.7 group was significantly lower than in the IgG2b isotype control group. In contrast, YUMM1.7, YUMM5.2, HCMel3, and B16-F10 tumors did not exhibit significant tumor growth or weight reduction in response to PD-L1 blockade (Fig. 1C). Thus, the YUMMER1.7 model was classified as ICB-sensitive, whereas YUMM1.7, YUMM5.2, HCMel3, and B16-F10 were considered ICB-resistant models.

Next, we assessed the occurrence of lymphatic metastasis by immunofluorescence staining of tdLN sections using melanoma markers in combination with CD45. Since the YUMM cell lines are amelanotic and lack expression of many classic melanoma markers, we used an antibody directed against the BRAF^V600E^ mutation, while the B16-F10 and HCMel3 tdLNs were stained with an antibody against gp100. Among the YUMM tumor models, YUMM5.2 showed the highest level of LN metastasis, with multiple metastatic foci spread over the whole LN (Fig. 1D, 1E), potentially explaining why only in this model mice had to be sacrificed due to weight loss before reaching the maximal tumor volume (Fig. 1C). YUMM1.7 tumors also metastasized spontaneously to LNs, although at a lower rate than YUMM5.2 cells. As expected, LN metastasis was not affected by PD-L1 blockade in these models. In the ICB-sensitive YUMMER1.7 model LN metastasis was barely detectable, even in control-treated mice. In contrast, gp100 staining in B16-F10- and HCMel3-draining LNs was scattered and may reflect a combination of tumor drainage and LN micrometastasis, as clearly stained gp100+ CD45-cell clusters were very rare. Nonetheless, B16-F10-draining LNs, in contrast to HCMel3-draining LNs, showed a small but significant decrease in gp100 staining after PD-L1 blockade (Fig. 1D, 1E). Furthermore, immunofluorescence analysis of primary tumor sections to detect lymphatic and blood vessels within the TME revealed a more extensive lymphatic and blood vessel network in YUMM compared to B16-F10 and HCMel3 tumors, but the vessel density was not affected by PD-L1 blockade (Fig. S1A, S1B).

### The CD8+ T cell profile is drastically changed after PD-L1 blockade in YUMMER1.7 primary tumors

To elucidate the immune landscapes and therapy responses associated with ICB-sensitive and -resistant melanoma, we harvested and processed both primary tumors and tdLNs for an in-depth comprehensive analysis of their immune landscape at the study endpoint. First, we globally assessed the frequency and status of the CD8+ T cell population using classic flow cytometry gating. Following PD-L1 blockade, we found that the frequency of CD8+ T cells decreased in B16-F10-draining LNs, while their frequency increased in the TME (Fig. 2A, 2B). In contrast, in the TME of ICB-sensitive YUMMER1.7, the frequency of CD8+ T cells surprisingly decreased after PD-L1 blockade. Notably, the exhaustion marker TOX was decreased in CD8+ T cells in both YUMMER1.7- and YUMM1.7-draining LNs (Fig. 2A, S2A), suggesting that ICB-mediated reduction of CD8+ T cell exhaustion in tdLNs is not necessarily associated with therapy success. In contrast, CD8+ TILs in the TME of only YUMMER1.7 melanoma exhibited a reduction in TOX level (Fig. 2B, S2A). Additionally, TCF1, a marker of naïve and stem-like T cells associated with a high proliferative capacity, specifically increased in the CD8+ T cells within the YUMMER1.7 TME (Fig. 2B, S2A). To analyze the CD8+ T cell space in greater detail, we used unsupervised clustering of cells stained with a multicolor flow cytometry panel including markers of CD8+ T cell subsets, activation and exhaustion states. We identified a total of 14 clusters of CD8+ T cells in tdLNs, comprising naïve, memory, tissue-resident, effector, progenitor-exhausted and exhausted subsets (Fig. S2B). Surprisingly, the CD8+ T cell subset landscape across the five melanoma-dLNs appeared very similar to that of naïve LNs, with naïve and CD103+ T cell clusters (clusters 5, 10, and 11) being the most abundant (Fig. S2C, S2D). Following PD-L1 blockade, we observed only minor changes in the frequency of CD8+ T cells per cluster among the 5 melanoma models, such as an increased population of granzyme B (GZMB)+ effector T cells (cluster 14) following ICB therapy in YUMMER1.7-draining LNs, with the highest frequency compared to the other models. Thus, CD8+ T cells in tdLNs are only minimally impacted by tumor growth and ICB therapy.

**Figure 2.**
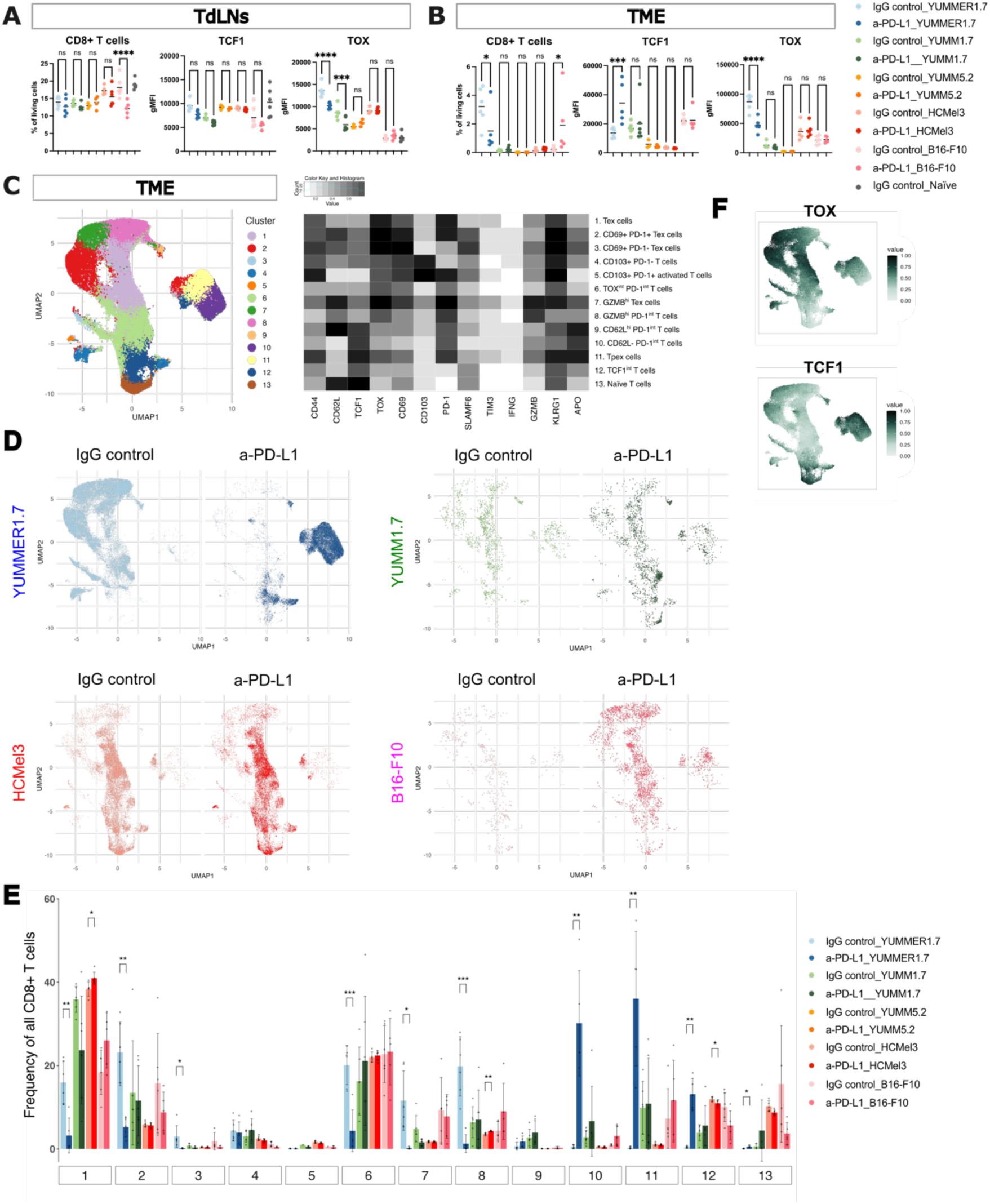
PD-L1 blockade affects the CD8+ T cell landscape only in the TME of sensitive tumors. **A-B**. Frequency of CD3+CD8+ cells and geometrical mean fluorescence intensity (gMFI) of TCF1 and TOX in the tdLNs (**A**) and TME (**B**) assessed by flow cytometry. N=3-6 mice/group. ** P < 0.01; *** P ≤ 0.001; **** P ≤ 0.0001; one-way ANOVA with Sidak post-test; ns=non-significant. **C.** UMAP with unsupervised clustering of CD3+CD8+ T cells in the TME and the respective heatmap depicting mean marker expression across all clusters. N=3-6 mice/group with 30-10.000 cells/mouse. **D.** Individual UMAP cluster plots of CD3+CD8+ T cells for each melanoma model and the indicated treatment. N=3-6 mice/group with 30-10.000 cells/mouse. **E.** CD3+CD8+ T cell cluster frequency in the TME. * P < 0.05; ** P < 0.01; *** P < 0.001; Student’s t-test. **F.** Projection of TCF1 and TOX transformed and normalized intensity on the clusters from (**C**). Tpex: progenitor exhausted T cell; Tex: exhausted T cell.

Within the TME, CD8+ T cells were most frequent in YUMMER1.7 and B16-F10 tumors, in comparison to the YUMM1.7, YUMM5.2, and HCMel3 models (Fig. 2B). Projection and clustering of the CD8+ T cells infiltrating primary tumors resulted in 13 distinct clusters (Fig. 2C). Due to the particularly low number of CD8+ T cells infiltrating YUMM5.2 tumors, we excluded this model from further analysis (Fig. 2B). In contrast to the LNs, CD8+ TILs in YUMMER1.7 tumors displayed a drastic remodeling after ICB treatment, indicating a massive impact of PD-L1 blockade especially on the TME (Fig. 2D, 2E). As expected from the manual gating (Fig. 2B), PD-L1 blockade increased TCF1 and decreased TOX expression in multiple T cell clusters (Fig. 2F). Furthermore, PD-L1 blockade increased the frequency of CD62L-PD-1^int^ T cells (cluster 10), Tpex (cluster 11), and TCF1^int^ T cells (cluster 12). In contrast, PD-1+ T cells (clusters 1, 2, 6, 7, and 8) and CD69+ PD-1-Tex cells (cluster 3) showed a drastic decrease (Fig. 2E).

Together, our data indicate that ICB therapy has only a minimal influence on CD8+ T cells in tdLNs which furthermore does not correlate with overall therapy responsiveness. On the other hand, ICB-induced invigoration of CD8+ T cells was prominent in the TME of ICB-sensitive melanoma but not in ICB-resistant melanoma.

### B cells do not respond to ICB therapy in melanoma

PD-L1 blockade was initially designed to reinvigorate T cells, but it is becoming increasingly clear that also other immune cell subsets express checkpoint receptors and might directly respond to ICB treatment, including B cells and innate, myeloid immune cells^23–25^. Therefore, we next assessed B cell (IgD+ B220+) subsets and responses to PD-L1 blockade in ICB-sensitive and -resistant melanoma-draining LNs. However, the frequency of these cells remained essentially unchanged (Fig. S3A). Unsupervised clustering revealed four main subsets of B cells, follicular B cells (cluster 8), plasma cells (cluster 7), and MHCII+ B cells (clusters 2 and 3) (Fig. S3B, S3C, S3D). ICB treatment had only minor effects on any of these. B cells in the TME were scarce, making it impossible to perform a subset analysis in this case. Thus, B cells appear to have limited relevance for melanoma immunity and ICB responsiveness.

### Distinct myeloid immune landscapes and ICB responses in melanoma-draining LNs and primary tumors

Finally, we turned to innate, myeloid immune cells, defined here as being CD11b+ and/or CD11c+. Using flow cytometry, we found that in most melanoma models, the overall frequencies of such cells in tdLNs and the TME were similar and did not change upon PD-L1 blockade (Fig. S4A, S4B, S4C). Only in B16-F10-draining LNs we noted a decrease in the frequency of myeloid cells. Interestingly, the frequency of myeloid cells infiltrating the YUMMER1.7 TME as well as classic activation markers related to antigen presentation and co-stimulation such as CD80, CD86, and MHC-II were decreased by PD-L1 blockade (Fig. S4A, S4C). Next, we used flow cytometry data projection and clustering analysis to define relevant myeloid subpopulations within tdLNs and primary tumors. Surprisingly, the myeloid immune landscape in tdLNs was very distinct in each of the models (Fig. 3A, 3B, 3C, S4D). Most strikingly, ICB-sensitive YUMMER1.7-draining LNs were populated by two clusters of CD11c^lo^ Ly6C^hi^ cells lacking classic markers of monocytes and macrophages (clusters 15 and 16), while YUMM1.7-draining LNs contained a high frequency of activated dendritic cells (clusters 7 and 11) and YUMM5.2-draining LNs a high frequency of granulocytes (cluster 4) (Fig. 3B, 3C, S4D). HCMel3-draining LNs showed a high presence of activated DCs (cluster 11), and B16-F10-draining LNs contained many Ly6C^hi^ F4/80^int^ cells (cluster 9) (Fig. 3C, S4D). Similarly, TME-infiltrating myeloid cell landscapes and ICB responses were clearly distinct between all the models (Fig. 3D, 3E, 3F, S4E). The YUMMER1.7 tumors were infiltrated by a high frequency of granulocytes (cluster 12), which further increased after PD-L1 blockade (Fig. 3E, 3F). On the other hand, two populations of tumor-associated macrophages (TAMs), one being high in MHCII suggesting a M1-like phenotype (clusters 4 and 7), were reduced in YUMMER1.7 tumors after ICB treatment (Fig. 3E, 3F). In contrast, ICB-resistant YUMM1.7 tumors were infiltrated by macrophages lacking MHC-II, suggesting an M2-like phenotype (cluster 8). Notably, in ICB-resistant melanoma models PD-L1 blockade had no obvious impact on myeloid infiltrates compared to the ICB-sensitive YUMMER1.7 model (Fig. 3E, 3F, S4E). Taken together, these results demonstrate that the myeloid immune landscape differs significantly among ICB-sensitive and -resistant melanoma, both within the TME and tdLNs. Furthermore, ICB therapy only impacted myeloid immune populations in the TME of sensitive melanoma.

**Figure 3.**
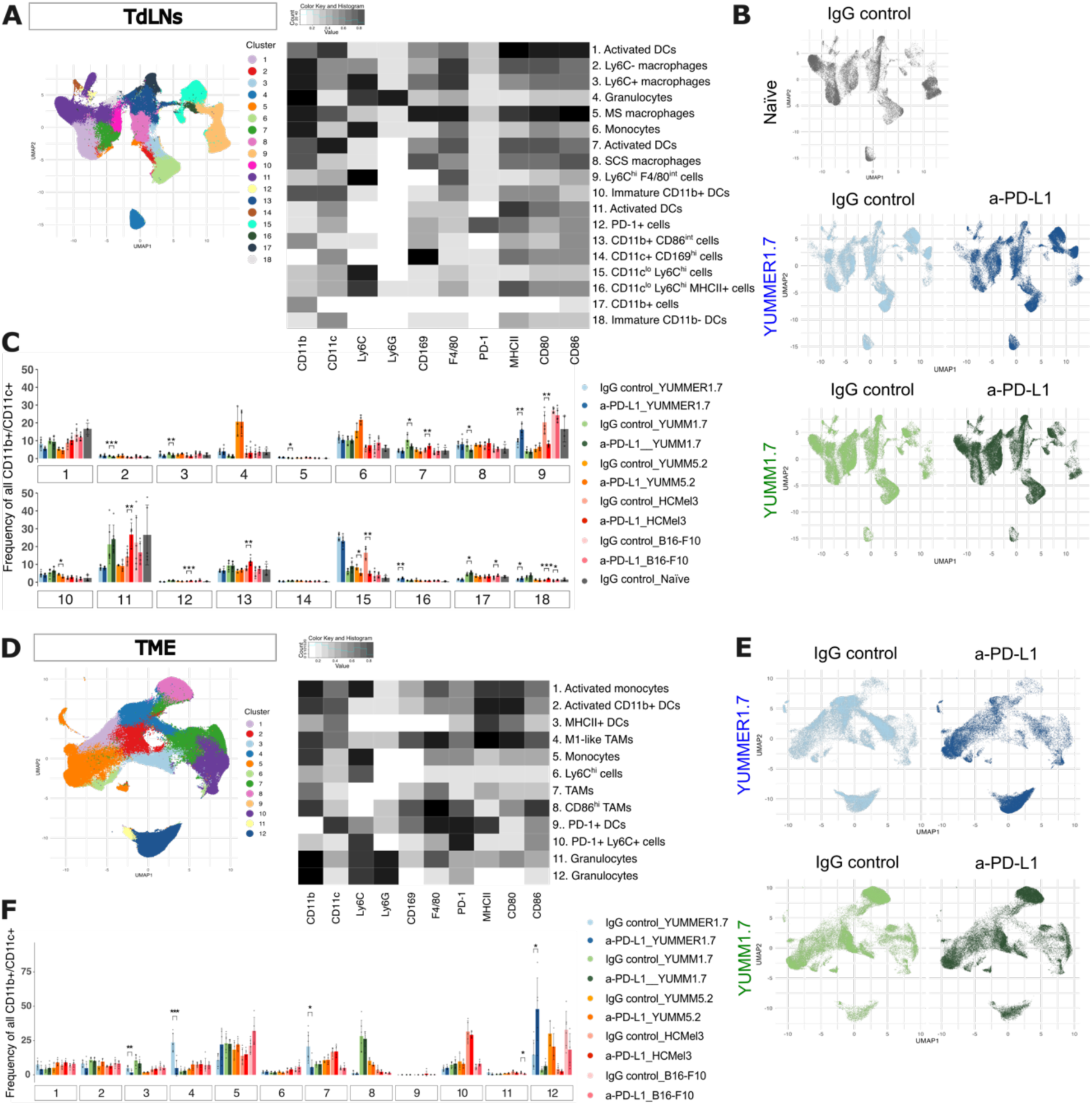
Myeloid immune landscapes and anti-PD-L1 responses in tdLNs and the TME diAer between melanoma models. **A.** UMAP and unsupervised clustering of CD11b+ and/or CD11c+ cells in naïve and tdLNs with heatmap representing the mean marker expression across all clusters. N=3-6 mice/group with 10.000 cells/mouse. **B.** Individual cluster plots of CD11b+ and/or CD11c+ cells in naïve, YUMMER1.7-, and YUMM1.7-draining LNs. N=3-6 mice/group with 10.000 cells/mouse. **C.** CD11b+ and/or CD11c+ cell cluster frequencies in naïve and melanoma-draining LNs. * P < 0.05; ** P < 0.01; *** P < 0.001; Student’s t-test. **D.** UMAP and unsupervised clustering of CD11b+ and/or CD11c+ cells in the TME with respective heatmap depicting mean marker expression across all clusters. N=3-6 mice/group with 530-10.000 cells/mouse. **E.** Individual cluster plots of the CD11b+ and/or CD11c+ cells infiltrating YUMMER1.7 and YUMM1.7 tumor under the indicated treatment. N=3-6 mice/group with 10.000 cells/mouse. **F.** CD11b+ and/or CD11c+ cell cluster frequency in the TME and the indicated treatment. * P < 0.05; ** P < 0.01; *** P < 0.001; Student’s t-test. DCs: dendritic cells; MS: medullary sinus; SCS: subcapsular sinus; TAM: tumor-associated macrophage.

### Single-cell RNA sequencing reveals myeloid subset identities in ICB-resistant and - sensitive melanoma

Strikingly different myeloid immune infiltrates in ICB-resistant vs. ICB-sensitive melanoma suggest that these infiltrates may have predictive potential for therapeutic outcomes. In particular, the prominent population of CD11c^lo^ Ly6C^hi^ cells (clusters 15 and 16) in YUMMER1.7-draining LNs caught our attention (Fig. 3C). However, as these cells lacked expression of classic monocyte, macrophage, DC and granulocyte markers, we aimed to further characterize them using single-cell RNA sequencing (scRNA-seq). To this end, we enriched CD11b+ and/or CD11c+ cells from LNs draining ICB-sensitive YUMMER1.7 tumors in comparison to the parental, ICB-resistant YUMM1.7 model as well as naïve LNs and performed scRNA-seq. After quality filtering, a total of 25.483 cells were subjected to unsupervised clustering and annotation. The cells clustered into 25 CD45 (*Ptprc*)-positive immune cell clusters (Fig. 4A) that could further be classified into groups of DCs, monocytes / macrophages, granulocytes, B-, T-, and innate lymphocytes cells (ILCs) based on established markers (Fig. S5A). In addition, we obtained a small cluster of CD45-negative endothelial cells (Fig. 4A, S5A).

**Figure 4.**
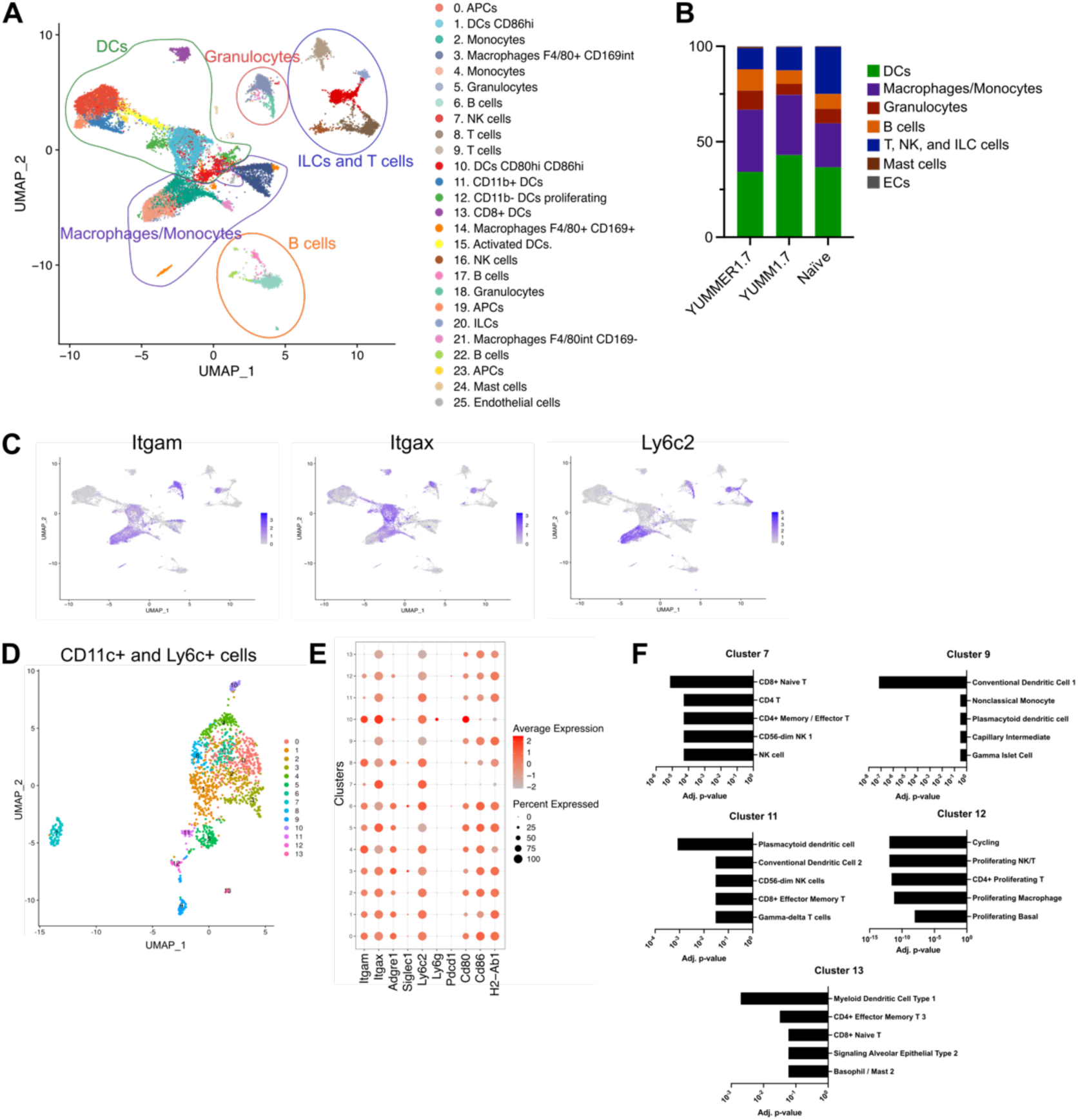
scRNA-seq analysis of CD11b+/CD11c+ cells identifies prominent myeloid immune subsets in ICB-sensitive melanoma-draining LNs. **A.** UMAP and unsupervised clustering of scRNA-seq data of CD11b+ and/or CD11c+ cells enriched from naïve, YUMMER1.7-, and YUMM1.7-draining LNs at day 14 post tumor inoculation, N=10889, 7656, 6936 cells, respectively. **B.** Frequency of major immune cell populations per model. **C.** Feature plots of selected marker genes *Itgam*, *Itgax*, and *Ly6c*2 projected onto the UMAP clusters from (**A**). **D**. UMAP and unsupervised re-clustering of cells expressing CD11c and Ly6C extracted from (**A**). **E**. Dot plot showing expression of all markers used for flow cytometry. **F**. Cell type prediction (Azimuth) for the cluster 7, 9, 11, 12 and 13, based on cluster-specific marker genes. DCs: dendritic cells; ECs: endothelial cells; APCs: antigen-presenting cells; pDCs: plasmacytoid dendritic cells.

In line with our flow cytometry-based analyses, the scRNA-seq data confirmed that YUMM1.7-draining LNs contained more DCs, including multiple distinct clusters representing various DC subsets and activation states (clusters 1, 10, 11, 12, 13, and 15), compared to YUMMER1.7-draining and naïve LNs (Fig. 4B, S5B, S5C). Differential gene expression analysis of an activated DC cluster (cluster 1) revealed that 70 genes were downregulated, and 148 genes were upregulated in YUMMER1.7-draining LNs compared to YUMM1.7-draining LNs, the latter comprising many genes related to type-I and -II interferon-mediated signaling (Fig. S5B, S5D, Supplementary Table 2). Additionally, YUMMER1.7-draining LNs exhibited a greater number of granulocytes (clusters 5 and 18) compared to the YUMM1.7-draining LNs, with 98 genes significantly upregulated in cluster 5 that were again associated with interferon responses (Fig. S5E, S5F, Supplementary Table 2). However, since the YUMM5.2 model displayed an even higher granulocyte population within tdLNs in our flow cytometry analysis (Fig. 3C), granulocytes are not a promising biomarker candidate for ICB responsiveness.

To learn more about the phenotype of the CD11c^lo^ Ly6C^hi^ cell populations enriched in ICB-sensitive YUMMER1.7-draining LNs, we closely inspected clusters with high Ly6C expression and low levels of CD11c and CD11b, in line with the flow cytometry data (Fig. 3A, 3B, 3C, 4C). However, we did not identify any single cluster matching this phenotype (Fig. S5G). Therefore, we extracted and re-clustered all cells expressing CD11c and Ly6C from our scRNA-seq dataset to analyze them more deeply (Fig. 4D). Notably, clusters 9, 11, 12, and 13 exhibited low level of CD11c and minimal CD11b expressions, but only cluster 11 showed high level of Ly6C, similar to cluster 15 and 16 from the flow cytometry data (Fig. 3C, 4E). Cluster 7 also exhibited high level of Ly6C and minimal level of CD11b but high level of CD11c. Interestingly, cell type prediction based on cluster-specific marker genes (Supplementary Table 3) strongly suggested that cluster 11 is composed of plasmacytoid dendritic cells (pDCs), whereas cluster 9 and 13 represent conventional DCs (cDCs), cluster 12 were proliferating cells, and cluster 7 were CD8+ T cells. (Fig. 4F).

### Plasmacytoid DCs are enriched in ICB-sensitive melanoma-draining LNs and correlate with good prognosis in melanoma patients

To confirm whether pDCs or cDCs are enriched in LNs draining ICB-sensitive YUMMER1.7 tumors, we analyzed LNs of an additional cohort of tumor-bearing mice with a modified myeloid flow cytometry panel which included the pDC marker Bst2. The panel also included markers for exclusion of lymphocytes and NK cells (CD3, CD19, and NK1.1) as these cells may express Ly6C upon stimulation in combination with low levels of CD11b or CD11c. We found that CD11c^lo^ Ly6C^hi^ Bst2^hi^ pDCs were significantly enriched in YUMMER1.7-draining LNs compared to YUMM1.7-draining LNs. In contrast, there were no differences in the frequency of cDCs (Fig. 5A). This increased frequency of pDC was probably dependent on factors drained from the TME, as non-involved popliteal LNs in tumor-bearing mice did not show any difference in the frequency of pDCs between the two models. PDCs made up a sizeable proportion of LN cells, more than what would have been expected from our scRNA-seq data. Probably this was due to suboptimal capture of pDCs by CD11c magnetic beads used for enrichment. Congruently, multiplexed immunofluorescence staining of tdLN sections further demonstrated that pDCs were readily detectable and primarily located near the medullary area of the LNs but could also be found scattered throughout the T cell zone in both tumor models (Fig. 5B).

**Figure 5.**
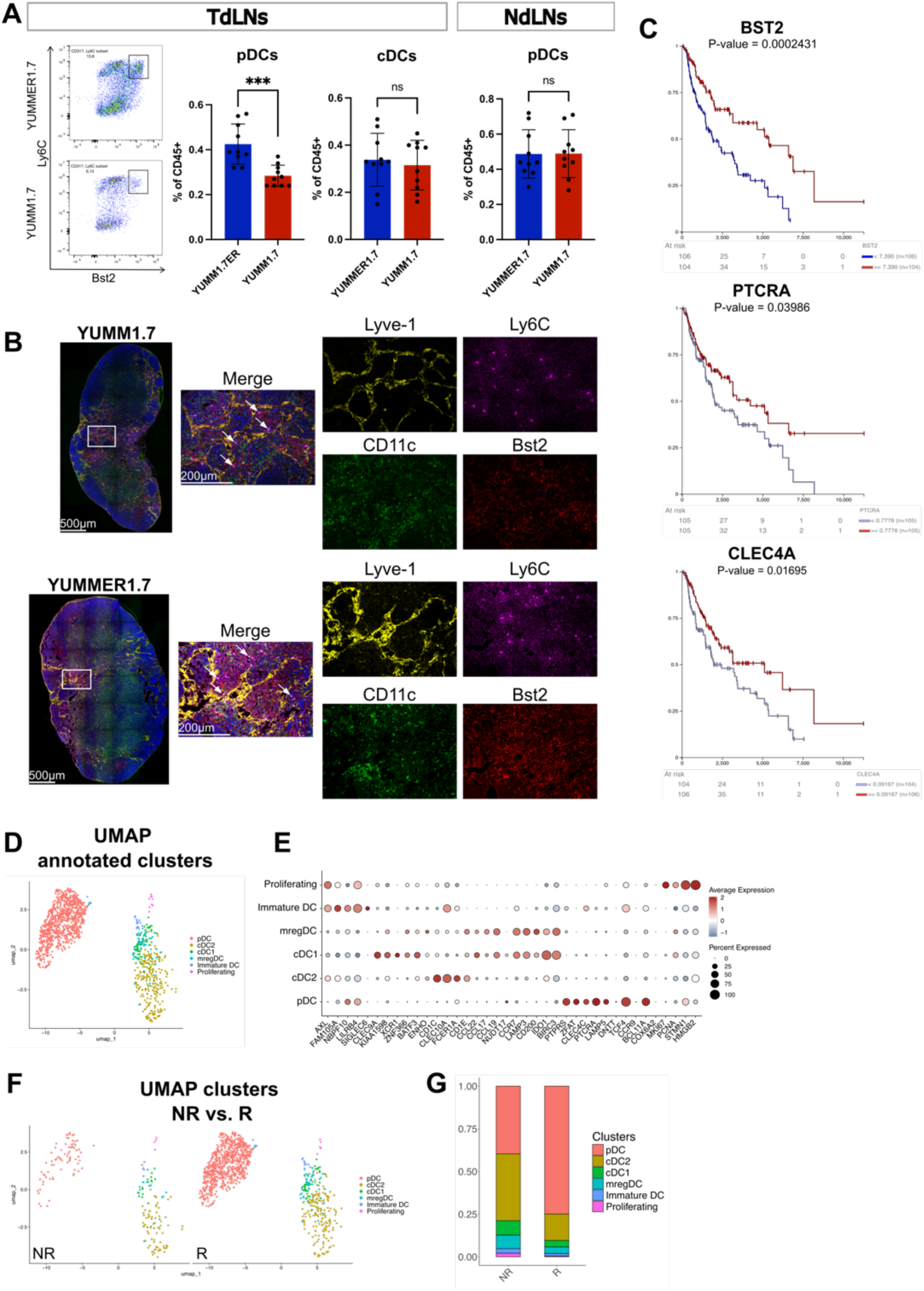
pDCs are enriched in ICB-responsive melanoma-draining LNs in mice and correlate with good outcome and ICB responsiveness in melanoma patients. **A.** Representative FACS plots and frequency of pDCs (CD11b-CD11c+ Ly6C+ Bst2+) and cDCs (CD11c+ MHCII^hi^) in YUMM1.7ER- and YUMM1.7-draining LNs and non-draining popliteal LNs of the same mice determined by flow cytometry. *** p < 0.001; Student’s t-test. **B.** Representative images of LN sections stained for CD11c (green), Ly6C (red), Bst2 (magenta), Lyve1 (yellow), and nuclear counterstaining (blue). **C**. Kaplan-Meier curves of selected pDC marker genes (*BST2, PTCRA, CLEC4A*) correlating with overall survival in human melanoma data (SKCM TCGA dataset, filtered for LN as site of resection / biopsy). The median value of gene expression was used as cutop. Statistical analysis was conducted using log-rank test. **D-E**. Unsupervised clustering (**D**) of DCs from metastatic melanoma LNs and dot plot (**E**) reflecting top marker gene expression used to discriminate the individual DC cluster from metastatic melanoma LNs. **F-G**. Comparison of DC clusters (**F**) in ICB-responders / tumor free patients (R) and -non-responders (NR) with the corresponding normalized frequency of cell clusters (**G**).

To determine whether the frequency of pDCs might be useful as predictive biomarker for ICB in melanoma patients, we examined the relationship between pDC marker gene expression and overall survival (OS) in the TCGA cutaneous melanoma (SKCM) cohort. Indeed, analysis of the dataset showed a significant positive correlation between high expression of pDC marker genes (*BST2, CLEC4C*, and *PTCRA*) with improved OS in patients for whom LN expression data was available (Fig. 5C). Additionally, we leveraged available scRNA-seq data of immune cells from metastatic LNs of melanoma patients subsequently responding to adjuvant ICB or remaining tumor free after surgery (responders, R) vs. patients progressing despite adjuvant ICB (non-responders, NR)^26^. By re-clustering of all cells classified as DCs, we found a total of 6 subsets, including pDCs (Fig. 5D, 5E). Notably, responders displayed a prominent pDC cluster within the DCs, in contrast to non-responders (Fig. 5F, 5G). Thus, our data suggests that the frequency of pDCs in tdLNs is a promising candidate marker to identify melanoma patients responding to ICB.

## Discussion

Although ICB has been used successfully for over a decade, our understanding of the precise mode of action and mechanisms underlying sensitivity and resistance towards this type of therapy is still limited. Originally designed to re-invigorate dysfunctional T cells within the TME, it has become clear that ICB outcomes are mediated by multilayered responses, including regional (i.e. in tdLNs) as well as systemic responses. Furthermore, the PD-1 / PD-L1 pathway, which is targeted by multiple clinically approved ICB agents, may be active in immune cell types other than T cells, including DCs and MDSCs^23,24^. Thus, at least in the context of PD-1 / PD-L1 blockade, direct effects on innate immunity may contribute to the efficiency of the therapy. At the same time, dissecting innate contributions to ICB efficiency may yield novel predictive biomarkers that are still urgently needed. In line with this, circulating innate immune cells such as monocytes and MDSCs have already been described to yield predictive power^27^.

Here, we comprehensively mapped immune landscapes of the TME and tdLNs in 5 syngeneic, orthotopically (intradermally) implanted melanoma models in mice, in the presence or absence of systemic ICB blocking PD-L1. Our flow cytometry analyses comprised panels designed to analyze subsets and activation states of T cells, B cells, and innate immune cells, combined with unsupervised clustering for unbiased subset identification. Of note, 4 of the 5 models were completely resistant to systemic PD-L1 blockade. Only the YUMMER1.7 model, derived from YUMM1.7 cells via somatic hypermutation^21^, showed decreased tumor growth upon PD-L1 blockade.

Our data clearly demonstrate that, whereas CD8+ T cells showed drastic remodeling with increased activation and reduced exhaustion patterns in the TME of the ICB-sensitive model in comparison to ICB-resistant models, CD8+ T cell landscapes in tdLNs were very similar in all the models and showed reduced exhaustion upon therapy in both ICB-sensitive and -resistant models (YUMMER1.7 and YUMM1.7, respectively). Thus, CD8+ T cell signatures in melanoma-draining LNs might not be useful to predict ICB efficiency. Similarly, B cells in tdLNs showed very similar profiles in ICB-resistant and -sensitive melanoma and were not significantly altered by systemic PD-L1 blockade either, suggesting that B cells are not directly relevant for the outcome of this type of therapy.

In contrast, we identified a striking divergence in innate myeloid immune profiles between ICB-sensitive and resistant melanoma, in both the TME and tdLNs. Surprisingly, ICB sensitivity was associated with a lower frequency of activated DCs. This might be due to the rapid killing of tumor antigen-presenting DCs by highly activated T cells. Importantly, we also noted a higher frequency of pDCs in LNs draining ICB-sensitive melanoma, and pDC frequency and marker gene expression correlated with good outcome and ICB responses in melanoma patients. Thus, our data suggests that assessment of pDC frequency in tdLN biopsies may be an impactful supplement to the currently applied predictive biomarkers.

pDCs are known to circulate in the blood and to populate secondary lymphatic organs such as LNs, where they predominantly reside in the medullary region. The most prominent function of pDCs is to act as rapid triggers of type-I interferon responses, releasing large amounts of interferon-alpha and -beta upon stimulation. Consequently, pDCs have been implicated in virus defense^28^. Additionally, pDCs may engage in antigen presentation to T cells, although they are considered less efficient than other DC subsets in this regard. Whether and how pDCs contribute to tumor immunity is so far poorly characterized. However, type-I interferons positively impact tumor immunity^29^. Congruently, we found that other myeloid immune cells in ICB-sensitive tdLN showed a gene expression profile indicative of type-I interferon responses. Furthermore, our finding that pDC marker genes correlate with favorable outcome in melanoma patients irrespectively of whether they were treated with ICB or not suggests that these cells may exert beneficial functions in tumor immunity.

Our study has certain limitations that require consideration while interpreting our results. On the one hand, most of the melanoma models used here, although well characterized and partly bearing clinically relevant mutations, are ICB-resistant, and only one model, YUMMER1.7, was sensitive to the therapy. Since YUMMER1.7 cells were generated through somatic hypermutation to overcome a lack of tumor-associated antigens recognizable by the immune system, our comparison may not be generalizable to other ICB-sensitive melanoma models. Furthermore, the translational evidence that pDCs may be a predictive biomarker for ICB in melanoma patients remains limited at this point, since the case numbers with available single-cell data derived from LNs is very small, and the analyzed cases with high pDC frequency were patients that had remained tumor free upon adjuvant ICB, not patients that had shown a clinical response to ICB^26^. Thus, in order to establish the power of a pDC-based predictive biomarker, larger patient cohorts need to be investigated retrospectively. Nonetheless, our results are promising and may spark renewed interest in an immune cell population long known for its role in viral defense but with still elusive implications for tumor immunity.

## Material and Methods

### Mice

Male C57Bl6/NRJ (referred to as C57BL/6) male mice were purchased from Janvier and housed at the Preclinical Models Core Facility at the Medical Faculty Mannheim. The mice had free access to food and drinking water and were utilized for tumor studies at an age of 8-10 weeks. All animal experiments in this study were approved by the responsible regional authorities (Regierungspräsidium Karlsruhe, permit G-177/22).

### Cell lines

YUMMER1.7 cells^21^ were kindly provided by Prof. Cornelia Halin (Swiss Federal Institute of Technology, Zurich), and the YUMM1.7 and YUMM5.2 cell lines^22^ were generously provided by Prof. Jonathan Sleeman (Medical Faculty Mannheim, Heidelberg University). The YUMM cell lines (YUMMER1.7, YUMM1.7, YUMM5.2) were cultured in DMEM/F12 media and supplemented with 1% non-essential amino acids (NEAA) and 10% fetal bovine serum (FBS). B16-F10^22^ cells were cultured in DMEM supplemented with pyruvate and glutamax, along with 10% FBS. HCMel3 cells^20^ were cultured in RPMI-1640 supplemented with 10% FBS, 1x L-glutamine, NEAA, 10 mmol/L HEPES, and 20 µmol/L β-mercaptoethanol (b-ME).

### Tumor models

One day prior to tumor injection, the lower back of C57BL/6 mice was shaved. Tumor cells were injected intradermally in both flanks. The following cell numbers were injected in a volume of 25 µL of PBS: 200,000 YUMM1.7 and YUMM5.2 cells, 300,000 YUMMER1.7 cells, 400,000 HCMel3 cells, or 500,000 B16-F10 cells. Mice were monitored three times per week, with tumor growth assessed through caliper measurements. When the tumor size reached a volume of 50 mm^3^, treatment with PD-L1 antibody (200 µg in 100 µL PBS, BioXcell, clone 10F.9G2) or an equal dose of IgG2b isotype control (BioXcell, clone LTF-2) was initiated via intraperitoneal injection and continued every third day for a maximum of four injections. Mice were euthanized when tumors reached a volume of 1 cm^3^ or the weight of the mice decreased by 20%.

### Tissue processing

Tumor-draining axillary and inguinal LNs, non-draining popliteal LNs as well as primary tumors from both flanks of the mice were harvested. One inguinal LN and half of the tumor tissue were cryopreserved for immunofluorescence analysis while the remaining samples were processed for flow cytometry analysis as described before^10^. In brief, the LN capsule was disrupted using a needle before being digested in a solution containing 1 mg/mL Collagenase type IV (Thermo) and 40 µg/mL DNase I (Roche) in DMEM supplemented with 2% FBS, 1.2 mM CaCl_2_ at 37°C for 20 minutes. After this initial digestion, the supernatant was collected, and the remaining LN fragments were digested using 3.5 mg/mL Collagenase type IV and 40 µg/mL DNase I in DMEM supplemented with 2% FCS, 1.2 mM CaCl_2_ at 37°C for 15 minutes. The cell suspension from each digestion was passed through a cell strainer, then resuspended in FACS buffer (1% FBS, 1 mmol/L EDTA, and 0.02% NaN_3_ in PBS) with anti-CD16/CD32 (BioLegend 101302) and kept on ice. For tumor processing, the tumors were cut into small pieces and directly digested with 3.5 mg/mL Collagenase type IV and 40 µg/mL DNase I in DMEM supplemented with 2% FCS, 1.2 mM CaCl_2_ at 37°C for 30 minutes. The digested tumors were then passed through a cell strainer. The resulting cell suspension was treated with an erythrocyte lysis buffer (0.15 mol/L NH_4_Cl, 0.01 mol/L KHCO_3_, 0.1 mmol/L Na_2_EDTA) before being resuspended in FACS buffer with anti-CD16/CD32 on ice.

### Flow Cytometry

Cell suspensions were stained with antibodies (see Supplementary Table 1) in 50uL of Brilliant stain buffer (BD) for an hour on ice. Then, cell suspensions were fixed and permeabilized overnight using the Foxp3 Transcription Factor Staining Buffer Set (Thermo) to stain with intracellular antibodies (see Supplementary Table 1). Flow cytometry data was acquired with a NothernLights instrument (Cytek) and analyzed using FlowJo v10.10.0 (BD).

High-parametric flow cytometry data projection and clustering were performed following the methods outlined by Ingelfinger et al.^31^. The unmixed flow cytometry data was compensated using FlowJo v10.10.0, and populations of interest (CD3+CD8+, IgD+B220+, and CD11b+ and/or CD11c+), were carefully gated. 10.000 cells per sample were exported whenever possible. If fewer than 10.000 cells were present, all available cells were exported and then imported into R (v4.4.2). The CyCombine package was utilized to combine the population of interest from the different models. After an arcsinh transformation and a normalization of the values between 0 and 1, the transformed and normalized data were projected using the Uniform Manifold Approximation and Projection (UMAP) algorithm. Unsupervised clustering was conducted using the FlowSOM algorithm, with the number of clusters selected based on the marker projection within the UMAP to ensure meaningful clustering. Plots were created using the ggplot2 package and statistical tests using Student’s t-test analysis.

### Immunofluorescence staining and analysis

LNs were fixed in 4% paraformaldehyde for 2 hours, washed once in PBS, and then incubated in 30% sucrose overnight before being cryopreserved in OCT compound (Tissue-Tek) and frozen at -80°C. Tumors were directly cryopreserved in OCT compound after harvesting and frozen at -80°C. Tissue sections of 7 μm were fixed with acetone and 80% methanol before being rehydrated in PBS. They were then incubated in a blocking solution containing 5% donkey serum, 1% bovine serum albumin (BSA), 0.05% NaN_3_, and 0.1% Triton X-100 in PBS for one hour. Next, tissue sections were incubated with primary antibodies in the blocking solution for another hour. The primary antibodies used included goat anti-Lyve1 (R&D AF2125), rat Meca-32 (BioLegend 120504), rabbit anti-BRAFV600E (Thermo MA5-24661), rabbit anti-gp100 (Abcam 137078), rat anti-CD45-biotin (BioLegend 103104), hamster anti-CD11c-FITC (BioLegend 117306), rat anti-Lyve1-eFluor570 (Thermo 41-0443-82), rat anti-Bst2-AF647 (BioLegend 127014) and rat anti-Ly6C-AF700 (BioLegend 128024). After washing the sections, they were incubated with appropriate secondary antibodies if needed (donkey anti-rabbit-AF594, donkey anti-goat-AF488, donkey anti-goat-AF594, donkey, anti-rat-AF594 (all from Thermo), streptavidin-AF488 (BioLegend 405235) and Hoechst33342 (Thermo)) in PBS for one hour. Finally, after washing again, the sections were mounted and scanned using an AXIO Scan.Z1 or Scan.7 (Zeiss), a Stellaris 8 confocal microscope (Leica) or a LSM800 confocal microscope (Zeiss).

Blind unmixing of the 5-color confocal images was performed with the LAS X 3D Viewer software (Leica) using automatic spectral unmixing with denoised background. Other images were analyzed using Fiji^32^. To quantify blood and lymphatic vascular density, a region of interest (ROI) containing tumor tissue was selected from the composite color image, excluding damaged areas, adjacent healthy tissues, and staining artifacts. The image was then split into separate channels corresponding to each antibody. After adjusting and applying an appropriate threshold for each channel, binary images were created. Particles above the threshold and > 20 µm², were analyzed to determine the number of vessels within the defined ROI.

To quantify BRAF+ metastatic areas in YUMM5.2, YUMM1.7, and YUMMER1.7 tdLNs, multiple sections from tumor-draining LNs separated by approximately 100 µm were stained for CD45 and BRAF^V600E^. Immunofluorescence channels were merged to identify CD45-/BRAF^V600E^+ cell clusters. Following a manual marking of the metastatic area, a particle analysis was performed, and the results were normalized to the dimensions of a pre-defined ROI.

Gp100 staining was used to determine the metastatic areas in HCMel3 and B16-F10-draining LN sections. After exclusion of CD45+ areas, a threshold was chosen manually for gp100, and a particle analysis was performed for objects > 10 µm^2^. The quantification results were normalized to the ROI.

### CD11b/CD11c+ cell enrichment and single-cell RNA sequencing

At the endpoint of tumor growth, tdLNs were harvested and weighed before being digested with 1 mg/mL Liberase DH (Roche) and 0.2 mg/mL DNase I (Roche) in DMEM, using a ratio of 1 mL per 50 mg of tissue. The digestion was carried out for 30 minutes at 37°C. Afterwards, the mixture was passed through a cell strainer and resuspended in FACS buffer. A small portion of the cell suspension was used for flow cytometry analysis, while the remainder was used to enrich myeloid cells using a combination of CD11b and CD11c magnetic beads (Miltenyi) as per the protocol.

Single-cell RNA sequencing (scRNA-seq) library preparation was carried out at the Next Generation Sequencing Core Facility at Medical Faculty Mannheim using the Chromium Next GEM Single Cell 39 protocol (v3.1, 10x Genomics). CDNA libraries were subjected to Illumina sequencing (paired-end, 150 bp) at BMKGene GmbH, Münster. CellRanger (v7.0, 10x Genomics) was used for sequencing data alignment and quantification. Subsequently, Seurat (v4.0.1) was employed for further analysis^33^. After filtering of dead, doublet, and outlier cells, 10889 naïve, 7656 YUMM1.7- and 6936 YUMMER1.7-derived cells were subjected to UMAP clustering and visualization. Differential gene expression analysis was conducted for clusters of interest by filtering for a g-value below 0.05. Gene Ontology (GO) analysis of differentially expressed genes was done using DAVID^34,35^. Cell type prediction of individual clusters was done using EnrichR^36^ based on significantly enriched marker genes.

### Human melanoma single-cell RNA sequencing

scRNA-seq data from human melanoma lymph nodes were obtained from Schlenker et al.^26^ and analyzed in R using the Seurat pipeline (version 5.0)^37^. Briefly, samples were merged and integrated using Harmony^38^. Initially, low-quality cells were filtered out, followed by count normalization, linear transformation with ScaleData, and dimensionality reduction using PCA. Graph-based clustering of lymph node cells was performed, enabling dendritic cell annotation based on cluster-specific genes identified through the FindAllMarkers function. Finally, the frequency of each dendritic cell cluster was calculated, stratifying patients into responders (R) and non-responders (NR) based on treatment outcome.

### TCGA data analysis

Correlation between selected genes and overall survival (OS) of melanoma patients (SKCM cohort) was investigated using the XENA browser^39^. The median expression level was used as cutoff to stratify into low- and high-expressing patients. Statistical analysis was done using a log-rank test.

### Statistics

GraphPad Prism (v10.2.3) and R (v4.4.2) were used to calculate the statistics. The statistical test analyses performed for each experiment is included in the figure legends. A p-value of less than 0.05 was considered statistically significant.

## Data availability

The single-cell RNA sequencing data reported in this manuscript will become accessible at GEO with accession number GSE293951 upon publication.

## Acknowledgement

The authors thank the SFB1366, the FlowCore, Preclinical Models, Next Generation Sequencing (NGS) and Live Cell Imaging (Lima) Core Facilities at the Medical Faculty Mannheim, Heidelberg University and the Leica Imaging Center at EMBL Heidelberg. The authors additionally acknowledge the data storage service SDS@hd supported by the Ministry of Science, Research and the Arts Baden-Württemberg (MWK) and the German Research Foundation (DFG) through grant INST 35/1503-1 FUGG, (Medical Faculty Mannheim, Heidelberg University). Furthermore, we thank Dr. Sparano and Dr. Mayoux (both Zurich University), Dr. Mahak Singhal, Dr. Chi-Chung Wu, Prof. Dr. Sleeman and Prof. Dr. Viktor Umansky (all Medical Faculty Mannheim, Heidelberg University) for technical support and fruitful discussions.

## Supplementary Figure Legends

**Supplementary Figure 1. Lymphatic and blood vascularization is not affected by PD-L1 blockade.**

**A.** Representative microscopy images of immunofluorescence staining for Lyve1 (green), Meca-32 (red), and DAPI (blue) of the indicated primary tumors treated with control IgG. **B.** Quantification of the lymphatic and blood vessel area and density within the primary tumor across the diperent melanoma models. ns=non-significant; one-way ANOVA with a Sidak post-test.

**Supplementary Figure 2. The CD8+ T cell landscape of tdLNs is minimally affected by PD-L1 blockade.**

**A.** Gating strategy used to identify CD3+CD8+ cells illustrated for a YUMMER1.7-dLN and primary tumor of a control IgG-treated mouse, including a representative histogram of TCF1 and TOX for the indicated treatment. **B.** UMAP and unsupervised clustering of CD3+CD8+ cells in naïve and tdLNs and the corresponding heatmap displaying the mean marker expression across all clusters. N=3-6 mice/group with 10.000 cells/mouse. **C.** Respective cluster plots of CD3+CD8+ T cells for naïve-and melanoma-draining LNs and the indicated treatment. N=3-6 mice/group with 10.000 cells/mouse. **D.** CD3+CD8+ T cell cluster frequency in tdLNs for the specified melanoma models and treatments. * P < 0.05; ** P < 0.01; Student’s t-test. Tpex: progenitor exhausted T cell; Tcm: central memory T cell.

**Supplementary Figure 3. PD-L1 blockade minimally impacts the B cell landscape in melanoma-draining LNs.**

**A.** Frequency of IgD+B220+ B cells in naïve and melanoma-draining LNs. ns=non-significant; one-way ANOVA with Sidak post-test. **B.** UMAP and unsupervised clustering of IgD+B220+ B cells in naïve and melanoma-dLNs with corresponding heatmap displaying the mean marker expression across individual clusters. N=3-6 mice/group with 10.000 cells/mouse. **C.** Individual UMAP plots of IgD+B220+ B cells in naïve and melanoma-draining LNs under the indicated treatment. N=3-6 mice/group with 10.000 cells/mouse. **D.** IgD+B220+ B cell cluster frequencies in naïve and melanoma-draining LNs under the indicated treatment. * P < 0.05; ** P < 0.01; *** P < 0.001; Student’s t-test. GC: germinal center.

**Supplementary Figure 4. Impact of PD-L1 blockade on myeloid immune cell phenotypes.**

**A.** Gating strategy used to identify CD11b+ and/or CD11c+ cells in tdLNs and primary tumors illustrated for a YUMMER1.7-dLN and primary tumor of an control IgG treated mouse, including a representative histogram of CD80, CD86, and MHCII for the indicated treatment. **B-C**. Frequency of CD11b+ and/or CD11+ cells and their geometrical mean fluorescence intensity (gMFI) of CD80, CD86, and MHCII in melanoma-dLNs (**B**) and the TME (**C**). N=3-6 mice/group. * P < 0.05; ** P < 0.01; ****, P ≤ 0.0001; one-way ANOVA with Sidak post-test. **D-E.** Individual UMAP plots of CD11b+ and/or CD11c+ cells in tdLNs (**D**) and the TME (**E**) in the YUMM5.2, HCMel3, and B16-F10 model. N=3-6 mice/group with 530-10.000 cells/mouse.

**Supplementary Figure 5. ScRNA-seq mapping of myeloid immune subsets in tdLNs in ICB-resistant and -sensitive melanoma.**

**A.** Feature plots for selected marker genes (*Ptprc, Ncr1, Cd3g, Ebf1, Ly6g, Csfr3, Itgam, Cx3cr1, Ccr7, Itgax, H2-Ab1* and *Adgre1*) illustrating the cluster annotation of the CD11b+ and/or CD11c+ cells from naïve, YUMMER1.7-, and YUMM1.7-draining LNs. **B.** Frequency of individual cell clusters from the scRNA-seq of CD11b+ and/or CD11c+ cells. **C.** Frequency of DCs (clusters 1, 10, 11, 12, 13, and 15) in naïve, YUMM1.7-, YUMMER1.7-draining LNs. **D.** Top 10 Gene Ontology of Biological Process (GO_BP direct) terms enriched in upregulated genes in activated DC cluster 1 in YUMMER1.7-compared to YUMM1.7-draining LNs. **E.** Frequency of granulocytes (clusters 5 and 18) in naïve, YUMM1.7- and YUMMER1.7-draining LNs. **F.** Top 10 GO_BP terms enriched among upregulated genes in granulocyte cluster 5 in YUMMER1.7-draining LNs. **G.** Dot plot showing expression of all markers used for flow cytometry across the scRNA-seq clusters.

**Supplementary Table 1. List of antibodies used for flow cytometry and immunostaining.**

**Supplementary Table 2. Differentially expressed genes in all scRNA-seq clusters.**

**Supplementary Table 3. Marker genes of the *Itgax*+ and *Ly6c2*+ subclusters.**

