## Supplementary figures and images for "Comprehensive immune profiling of melanoma-draining lymph nodes identifies plasmacytoid dendritic cells as new biomarker for PD-L1 blockade"

### Figure S1

#### Supplementary figure 1.

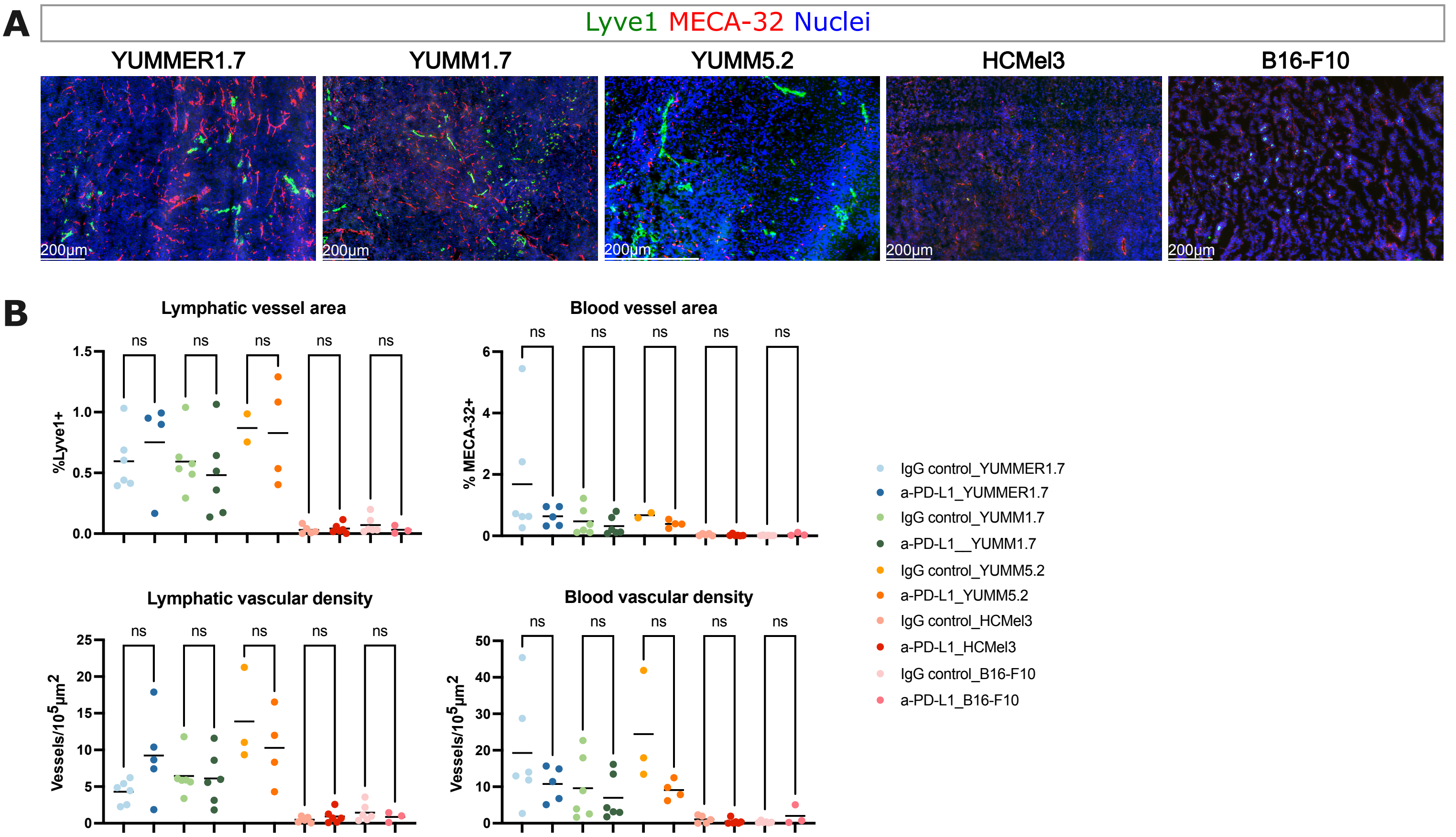

### Figure S2

##### Supplementary figure 2.

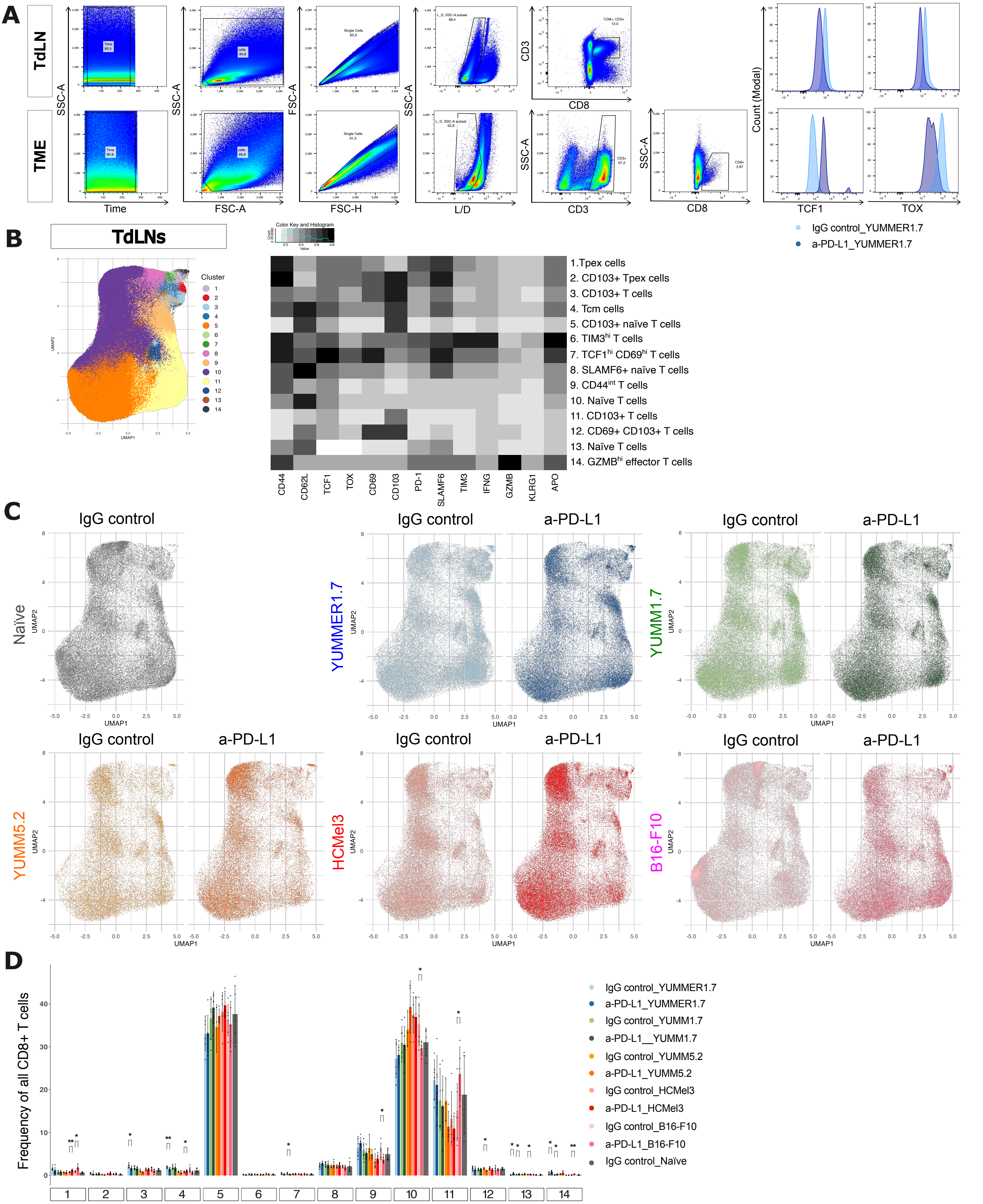

### Figure S3

#### Supplementary figure 3.

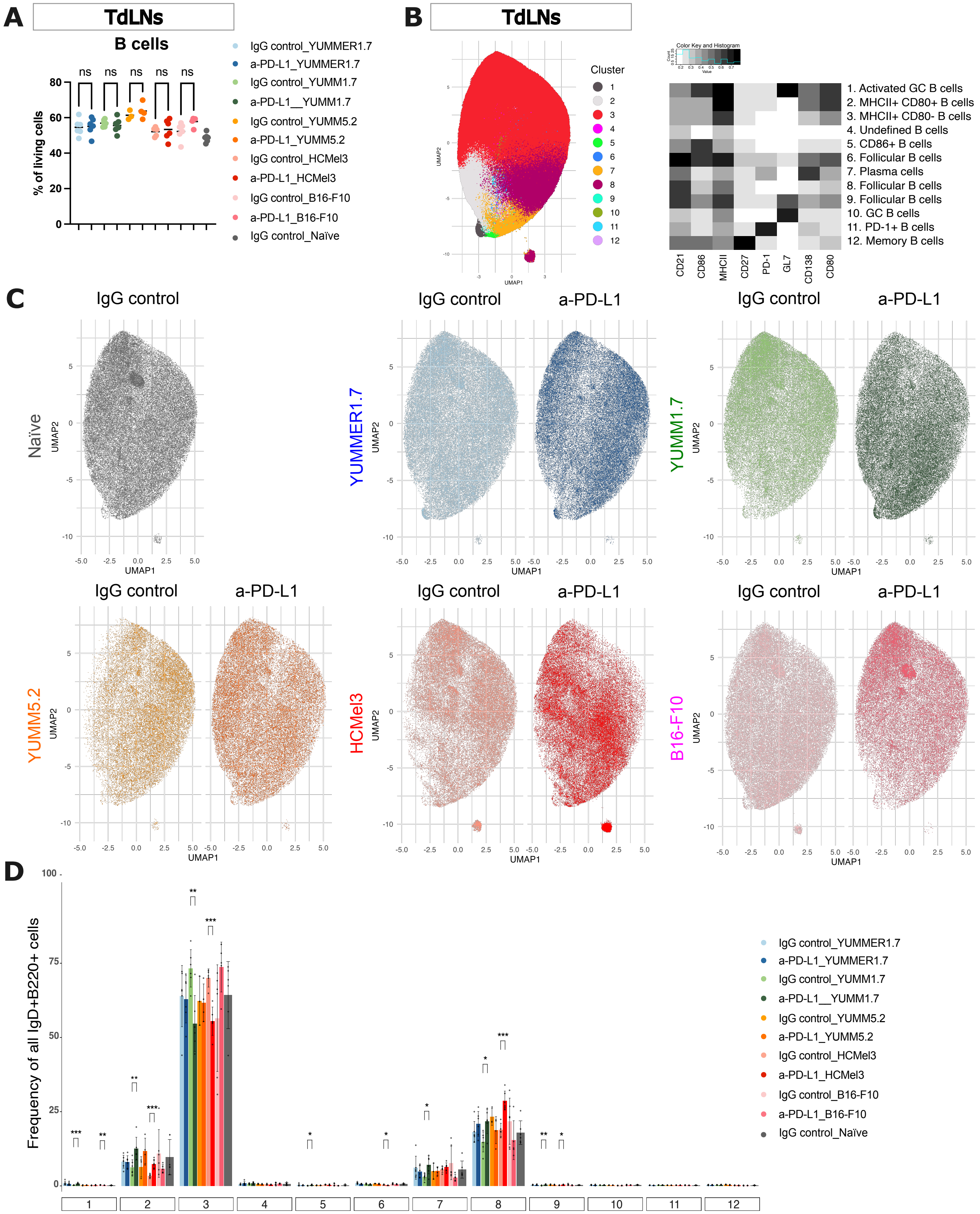

### Figure S4

##### Supplementary figure 4.

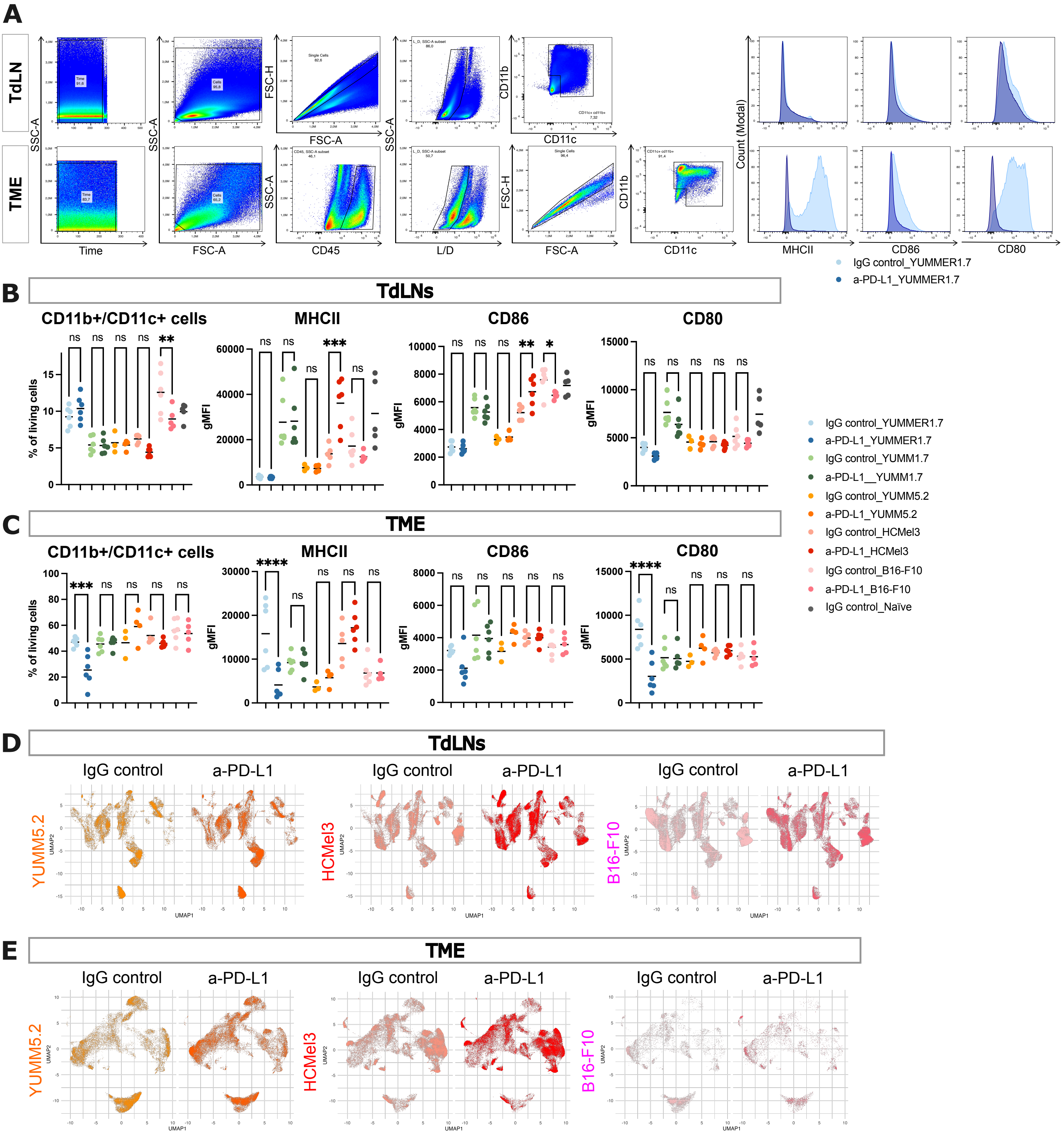

### Figure S5

#### Supplementary figure 5.

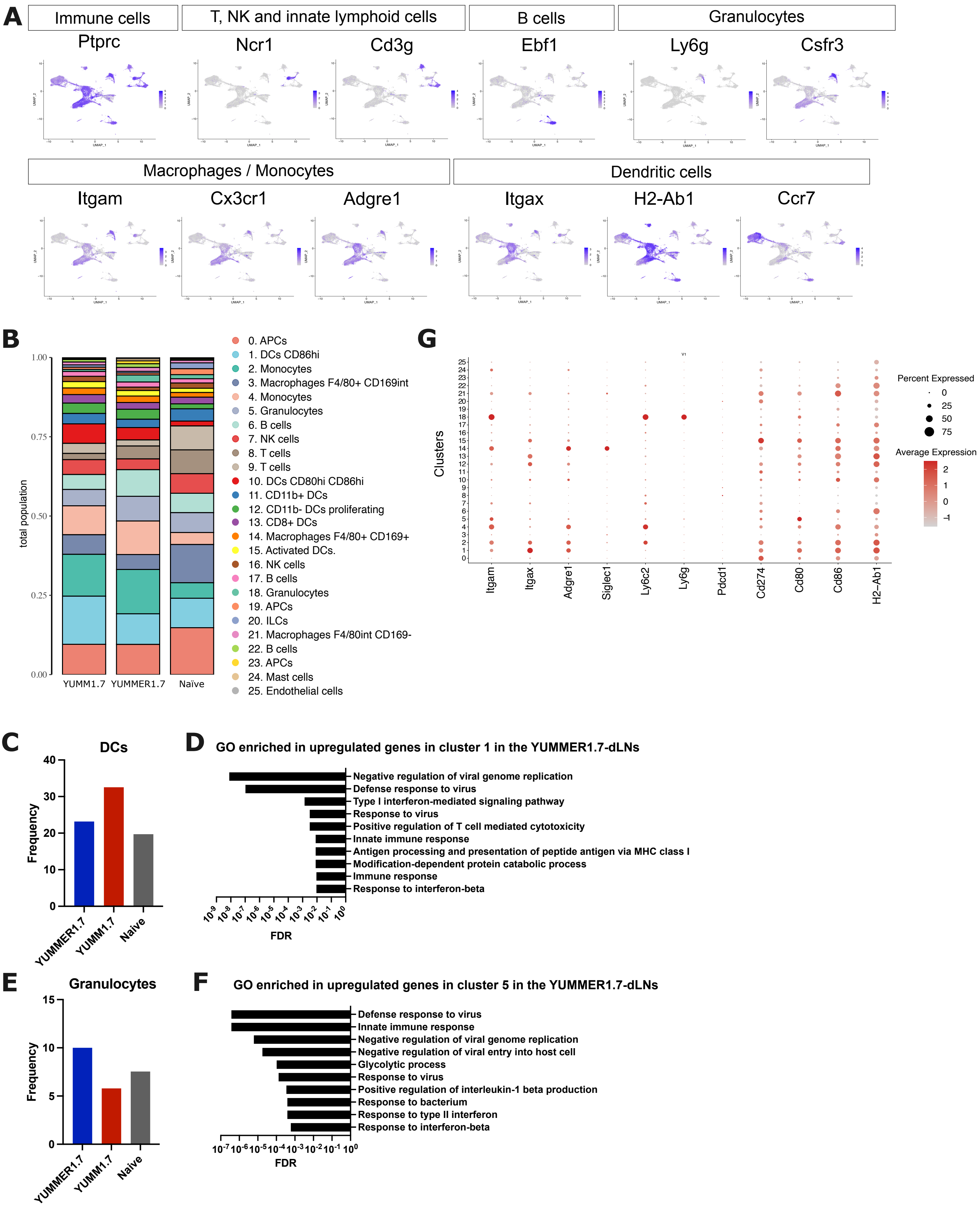
